# The challenge of distinguishing cell-cell complexes from singlet cells in non-imaging flow cytometry and single-cell sorting

**DOI:** 10.1101/2020.01.30.927137

**Authors:** Julie G Burel, Mikhail Pomaznoy, Cecilia S Lindestam Arlehamn, Gregory Seumois, Pandurangan Vijayanand, Alessandro Sette, Bjoern Peters

**Author notes:** Correspondence to: Dr Julie Burel, La Jolla Institute for Immunology, 9420 Athena Circle, La Jolla, CA 92037, USA.

## Abstract

Our recent work has highlighted that care needs to be taken when interpreting single cell data originating from flow cytometry acquisition or cell sorting: We found that doublets of T cells bound to other immune cells are often present in the live singlet gate of human peripheral blood samples acquired by flow cytometry. This hidden ‘contamination’ generates atypical gene signatures of mixed cell lineage in what is assumed to be single cells, which can lead to data misinterpretation, such as the description of novel immune cell types. Here, based on the example of T cell-monocyte complexes, we identify experimental and data analysis strategies to help distinguishing between singlets and cell-cell complexes in non-imaging flow cytometry and single-cell sorting. We found robust molecular signatures in both T cell-monocyte and T cell-B cell complexes that can distinguish them from singlets at both protein and mRNA levels. Imaging flow cytometry with appropriate gating strategy (matching the one used in cell sorting) and direct microscopy imaging after cell sorting were the two methods of choice to detect the presence of cell-cell complexes in suspicious dual-expressing cells. We finally applied this knowledge to highlight the likely presence of T cell-B cell complexes in a recently published dataset describing a novel cell population with mixed T cell and B cell lineage properties.

## Introduction

Multiparametric flow cytometry is a powerful tool to unravel the phenotypic heterogeneity of immune cells in humans. When combined with cell sorting and sequencing, it can unravel both protein and RNA expression programs within cell populations, which has led to the discovery of many novel immune cell subsets and associated functions, in both healthy and disease settings (1). However, our recent work highlights an occasionally underappreciated challenge, namely that care needs to be taken when interpreting single-cell data originating from flow cytometry acquisition or cell sorting: We found that when analyzing human peripheral blood monocular cells (PBMC), a small but reproducible proportion of presumed singlets by flow cytometry are tightly bound cell-cell complexes. These ‘contaminating’ dual cell complexes can mislead subsequent interpretation of what is presumed to be single-cell data.

We first identified by flow cytometry a cell population in the live singlet gate of human PBMC from patients with dual expression for CD3 and CD14 (2). We found that the frequency of these CD3+CD14+ cells was modulated as a result of immune perturbations such as vaccination, disease treatment and disease severity. This cell population expressed pan-markers of monocytes and T cells, both at the protein and the mRNA level, initially suggesting the discovery of a novel cell type with both T cell and monocyte lineage properties. However, subsequent analyses revealed that the CD3+CD14+ cell population did not consist of single cells bearing both T cell and monocyte lineage markers, but were either dual cell-cell complexes of T cells and monocytes, or T cells bound to smaller particles containing monocyte markers. Neither of these types of complexes were removed by conventional forward and side scatter gating approaches to avoid cell aggregates in flow cytometry. Importantly, the T cell-monocyte complexes we detected showed LFA-1/ICAM-1 polarization at their point of contact, could be isolated from fresh PBMC and whole blood, and were stable over time within a given individual, suggesting their presence *in vivo*, and not resulting from *ex vivo* sample manipulation. This makes studying the presence and composition of these complexes biologically important, and clearly refute our first interpretation that CD3+CD14+ cells could represent a novel cell type with mixed lineage properties.

Since T cell interactions are not restricted to monocytes, we expect to find them in complexes with other types of antigen-presenting cells (APCs). Indeed, others have previously reported on CD3+CD20+ cells detected by flow cytometry in human peripheral blood as doublets of T cells and B cells (3). Moreover, CD4+CD19+ cells have also been detected in draining lymph nodes of mice following infection, and found to be complexes of T follicular helper (Tfh) cells and B cells (4). Strikingly, the polarization of the Tfh cell and the immunoglobulin isotype class switch in the B cell were matching in each conjugate, and B cells in the conjugates were associated with a greater number of somatic hypermutations compared to singlets. Taken together, these results support our hypothesis that functional complexes of T cells and B cells are present *in vivo* and can be detected *ex vivo* by flow cytometry.

Here, we investigate how the presence of T cell-APC complexes in presumed single cell populations in flow cytometry can impact both data analysis and interpretation. We also highlight several experimental and data analysis strategies that can help identify complexes and thus avoid misinterpretations. We finally apply this strategy to follow up on a recently published study describing a novel immune cell population with T cell-B cell mixed lineage properties.

## Results

First, we assessed the impact of the presence of complexes on the resulting protein and gene expression profiling data generated with single cell techniques such as flow cytometry and single-cell sort RNA sequencing. We took as an example CD3+CD14+ cells, since we have already validated in our previous study that this cell population mainly consist of T cell-monocyte complexes and not CD3/CD14 dual expressers. Expression of T cell and monocyte markers in CD3+CD14+ cells was compared to singlets monocytes and T cells (see gating strategy **Figure 1A**) at both protein and mRNA level. Since T cell and monocyte markers are exclusively expressed by each cell type, when analyzed by flow cytometry, the CD3+CD14+ population shows similar expression levels of canonical markers for T cells and monocytes when compared to their singlet counterpart (**Figure 1B**). Similarly, at the mRNA level, when we performed single-cell sorts of the CD3+CD14+ population and subsequent RNA sequencing analysis of these cells, they showed positive expression for both T cell and monocyte markers (**Figure 1C**) and in PCA they represented a ‘halfway’ population between T cells and monocytes (**Figure 1D**). This reflects the simultaneous detection of the RNA content of both the T cell and the monocyte present in each sorted complex. Taken together, solely based on typical protein and gene expression profiles, complexes that are present within the singlet gate population in flow cytometry will appear like a composite cell population combining features of its two cell type components, similar to what a dual expresser would look like.

**Figure 1:**
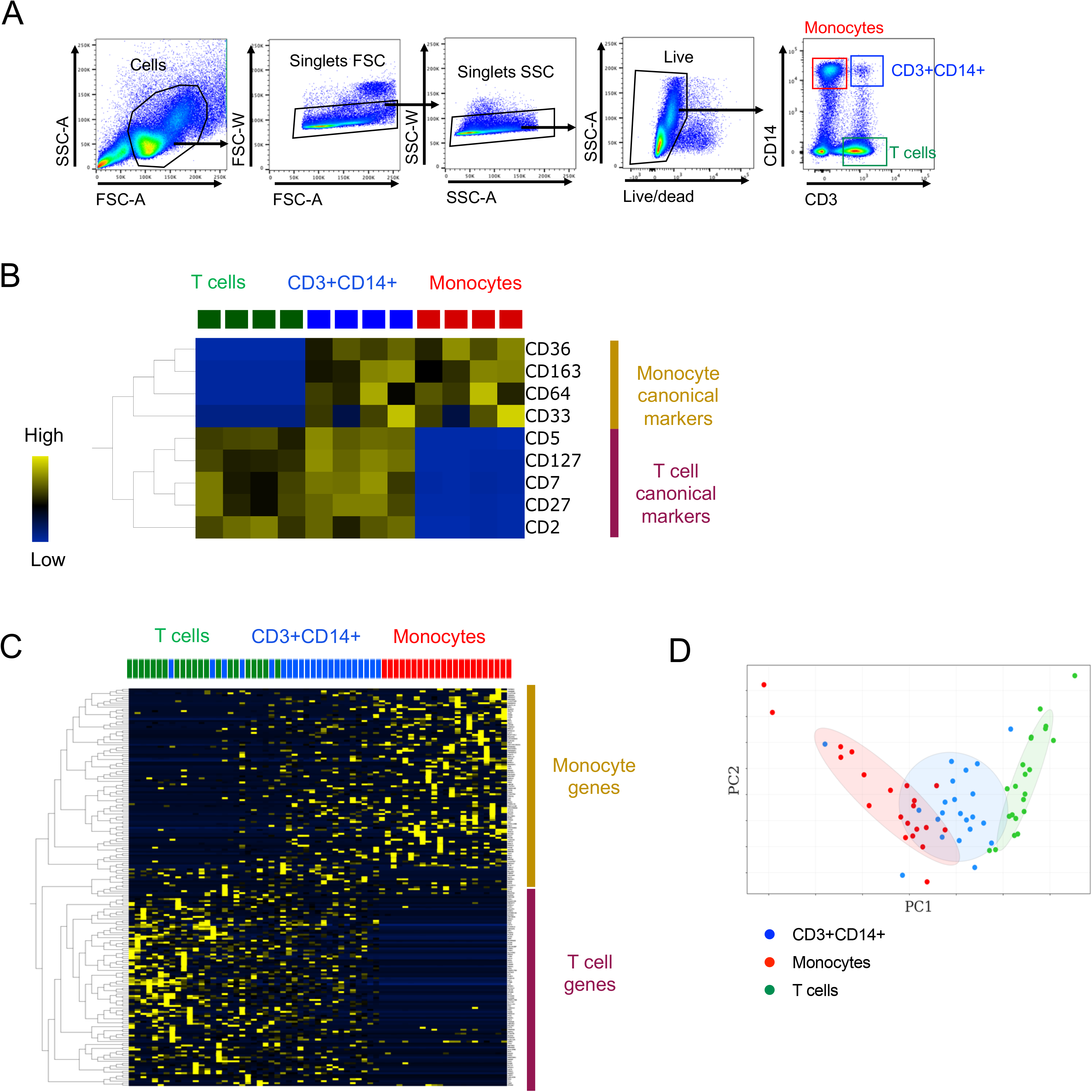
Cell-cell complexes appear like a ‘halfway’ cell population with mixed lineage properties from its two cell type components. A) Gating strategy to identify T cells, B cells and CD3+CD14+ cells in human PBMC. B) heatmap representing the protein expression of T cell and monocyte canonical markers in T cells, monocytes and CD3+CD14+ T cells. Expression was measured by non-imaging flow cytometry and expressed as median fluorescent intensity. C) Heatmap representing single cell gene expression of the 100 genes most upregulated in monocytes vs T cells and vice versa for sorted monocytes, T cells and CD3+CD14+ cells. D) PCA on all genes from the scRNAseq data of all three sorted populations. T cells, monocytes and CD3+CD14+ cells were identified as represented in Figure 1A. (A-B) Data were derived from PBMC of 4 healthy subjects. (C-D) Data were derived from 21 monocytes, 22 T cells and 22 CD3+CD14+ cells isolated from PBMC of one healthy subject, and are available under GEO accession number GSE117435.

Next, we looked for additional parameters in data generated from flow cytometry or single-cell RNAseq (scRNAseq) that could differentiate complexes from singlets. We have found it overall challenging to differentiate between T cell-monocyte complexes from singlets using non-imaging flow cytometry. Nevertheless, our initial study identified a couple of parameters that could partially distinguish doublets from singlets, even within the live singlet population (2). In flow cytometry, whereas FSC-A vs SSC-A gating cannot completely separate CD3+CD14+ cells from bigger singlet cells such as monocytes, median fluorescence intensity (MFI) for FSC-A or SSC-A is slightly increased for T cell-monocyte complexes when compared to singlet T cells or monocytes (**Figure 2A**). Thus, a stringent FSC-A vs SSC-A gating on lymphocytes such as T cells will remove most of cell-cell complexes, but will also exclude bigger lymphocytes such as activated T cells, and most of the monocyte population (**Figure 2A**). Similarly, staining for CD45 is increased in T cell-monocyte complexes compared to singlets (**Figure 2B**), as both cells express this marker. In RNAseq, TPM-transformed expression values identify the relative concentration of different RNA species rather than providing an absolute quantification (such as in flow cytometry). Thus, in scRNAseq, markers that are exclusively found in T cells or exclusively found in monocytes are about half that expression level when evaluating complexes compared to singlet T cells and monocytes (**Figure 2C**). Whereas these parameters are not sufficient to completely separate cell-cell complexes from single cells, they together represent useful signatures at both protein and mRNA level that can be checked by investigators after the data has been generated, and prompt for further cell-cell complexes analyses.

**Figure 2:**
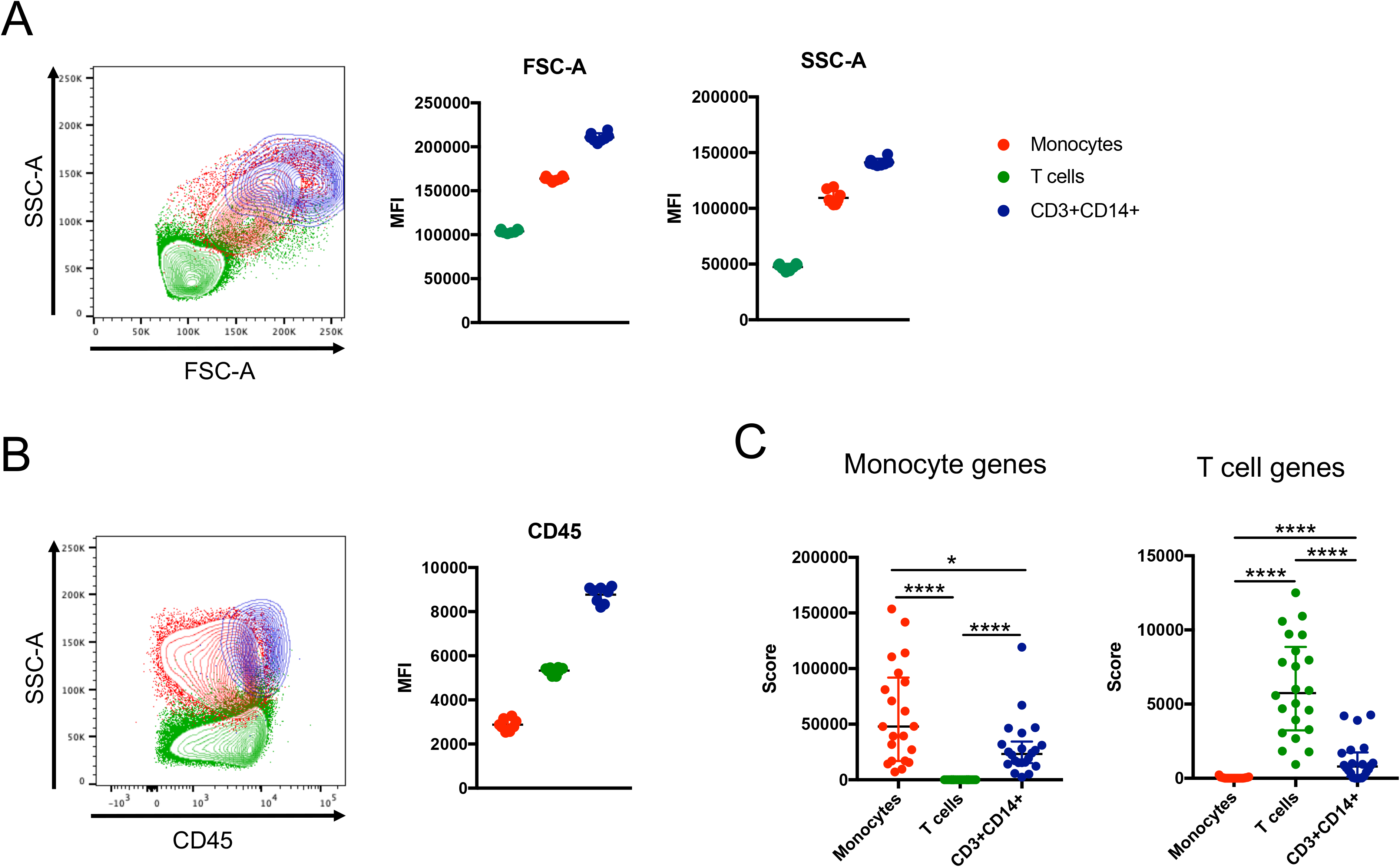
Molecular signatures of cell-cell complexes in flow cytometry and single-cell RNAseq derived from cell sorting. 2D representation plots and median fluorescence intensity (MFI) of A) FSC-A and SSC-A parameters and B) CD45 expression in monocytes, T cells and CD3+CD14+ cells in non-imaging flow cytometry. Data were derived from PBMC of 8 healthy subjects. C) Monocyte and T cell genes scores derived from the scRNAseq data of sorted monocytes, T cells and CD3+CD14+ cells as identified in Figure 1A. Data were derived from 21 monocytes, 22 T cells and 22 CD3+CD14+ cells isolated from PBMC of one healthy subject, and are available under GEO accession number GSE117435.

The by far preferred method to identify the presence of cell-cell complexes in a previously uncharacterized cell population is to use imaging flow cytometry. The typical gating strategy is, similarly to non-imaging flow cytometry (**Figure 1A**): First identify cells, followed by dead cells and doublets exclusion, to finally identify the double positive population (optimal (OPT) imaging gating, **Figure 3A**). However, imaging flow cytometry can identify doublets in a much more dispositive fashion than a standard flow cytometer by generating metrics derived from an actual picture of the event detected, which allows for the calculation of brightfield area and aspect ratio parameters. Additionally, the accuracy of the doublet exclusion gating can be cross-checked by viewing an image gallery of all gated objects. When we applied the OPT imaging gating strategy to the detection of CD3+CD14+ cells, it detected a much lower frequency of double positive events compared to non-imaging flow cytometry (**Figure 3C**). This signifies that the doublet exclusion in the imaging flow cytometer is far more sensitive than what we can expect from cell sorters relying on FSC and SSC based doublet exclusion, and results from the two cannot be directly compared. The first gating step for cell selection in imaging flow cytometry (SSC-A vs Brightfield Area) is sufficient to eliminate bigger aggregates (but not tight doublets) in a similar fashion that A vs. W, W vs. H or A vs. H would perform in non-imaging flow cytometry (**Figure 3B**). Using this cell sorting matching (CSM) gating strategy, we identified frequencies of CD3+CD14+ cells similar to what cell sorters detected (**Figure 3C**). Importantly, when exporting a gallery of events for both gating strategies, we identified that the CSM gating strategy identified mostly T cell-monocyte complexes (**Figure 3D**), whereas the OPT imaging gating strategy identified predominantly dual expressers (**Figure 3E**). Thus, the CSM gating strategy is more closely mimicking the population detected by cell sorters and non-imaging flow analyzers, and is a more reliable way to identify the possible presence of cell complexes and their relative proportion over true dual expressers within an unknown dual-positive cell population. In the case of CD3+CD14+ cells, this experiment demonstrates that whereas dual expressers do exist, the vast majority of dual positive events identified by flow cytometry and cell sorting represent T cell-monocyte complexes.

**Figure 3:**
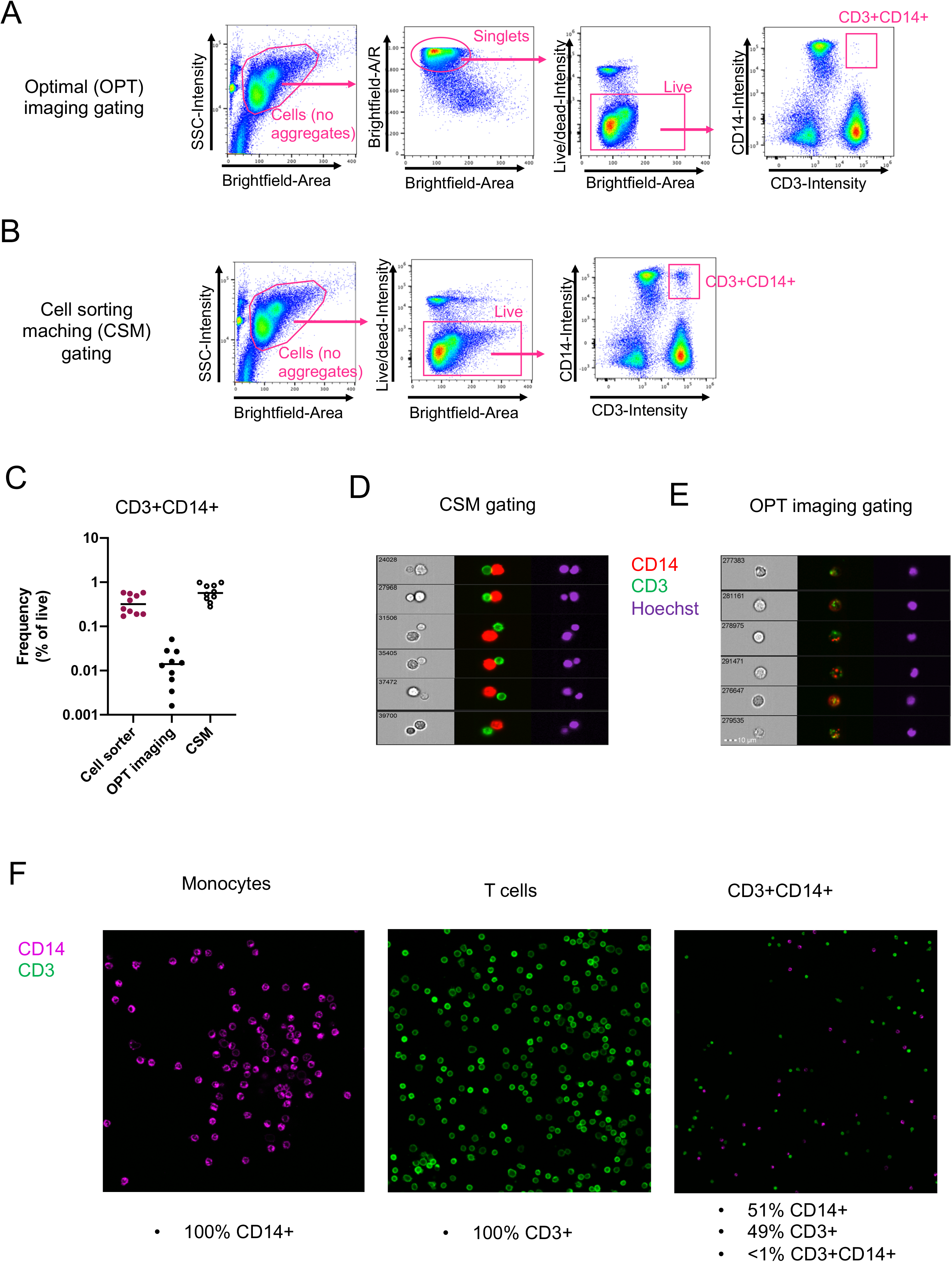
Imaging analyses to investigate the presence of cell-cell complexes in suspicious dual-expressing cell populations identified by non-imaging flow cytometry. A) Optimal (OPT) imaging and B) cell sorting matching (CSM) gating strategies to identify CD3+CD14+ cells with imaging flow cytometry. C) Frequencies of CD3+CD14+ cells detected by imaging flow cytometry using the OPT imaging vs the CSM gating strategies, in comparison to cell sorting. Random galleries of CD3+CD14+ events identified by imaging flow cytometry with D) the CSM gating strategy and E) the OPT imaging gating strategy. F) Confocal microscopy analysis of sorted monocytes, T cells and CD3+CD14+ cells as gated in Figure 1A. Data were derived from PBMC of 10 healthy subjects.

To circumvent the hurdles associated with matching gating strategies between imaging flow cytometry and cell sorting aforementioned, an alternative experiment is to directly sort the dual-expressing cell population onto a slide and analyze them using microscopy. In the case of CD3+CD14+ cells, we observed that cells identified as dual expressers by FACS represented a perfect stoichiometric mix of individual T cells and monocytes, but not dual expressers under the microscope (**Figure 3F**).

Finally, we applied our expertise in T cell-monocyte complexes detection to other types of cell-cell complexes that can be present in PBMC. A recent paper published in Cell (5) describes the discovery of a novel immune cell type expressing both T cell and B cell receptors in the peripheral blood of type I diabetes patients. These findings reminded us of our initial assumption that we had identified a novel cell population with both T cell and monocyte lineage properties. Thus, we next wondered whether the dual expressers (DE) cells identified by Ahmed al. indeed represented a completely novel population of a combined T cell and B cell lineage, or if they might be impacted by complexes of T cells and B cells tightly bound together that are circulating *in vivo*.

Following the same gating strategy as in the Ahmed et al. study, we were successfully able to detect DE cells (CD5+CD19+TCRab+ population within the live singlet gate) in human PBMC from healthy subjects (**Figure 4A**). Like CD3+CD14+ cells, DE cells were on average associated with higher FSC/SSC values, as well as higher CD45 staining intensity compared to T cells, B cells and CD5+CD19+TCRab-cells (**Figure 4B-C**). Imaging flow cytometry using the CSM gating strategy identified DE cells at a similar frequency traditional flow cytometry (**Figure 4D**), and revealed that DE cells predominantly consist of T cell-B cell complexes (**Figure 4E**). Contrastingly, the OPT imaging gating strategy identified a much lower frequency of double positive events (**Figure 4D**), all of which were dual expressers (**Figure 4F**). Thus, in our hands, DE cells in human PBMC display imaging and non-imaging flow cytometry characteristics of dual cell-cell complexes similar to what we found for T cell-monocyte complexes.

**Figure 4:**
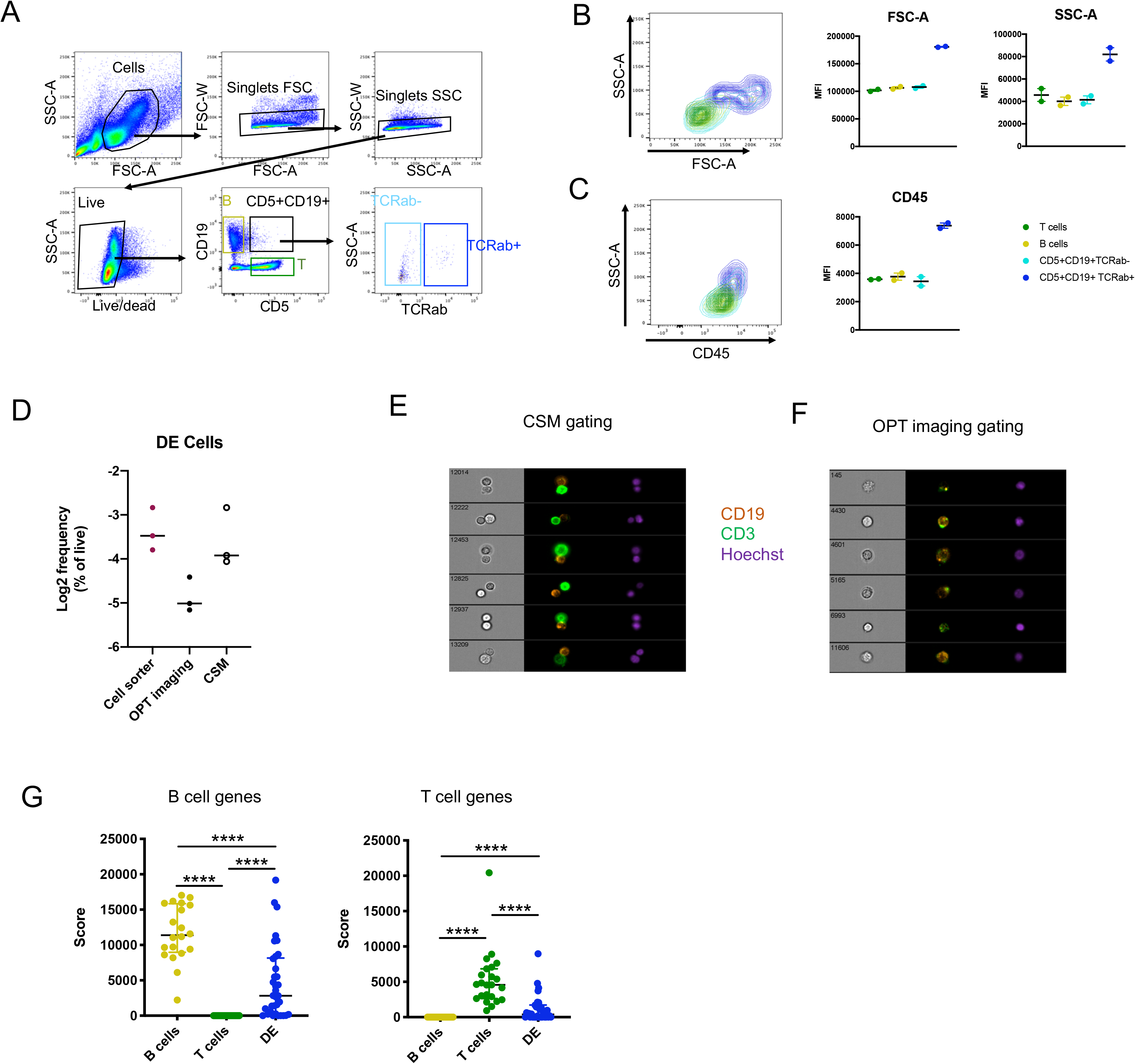
DE cells (CD5+CD9+TCRab+) in human PBMC have a flow cytometry and single-cell RNAseq signature of cell-cell complexes. A) Gating strategy to identify DE cells (CD5+CD19+TCRab+), CD5+CD19+TCRab-cells, T cells and B cells in human PBMC. Representative contour plot and median fluorescent intensity (MFI) of B) FSC-A vs SSC-A and C) CD45 in DE cells, CD5+CD19+TCRab-cells, T cells and B cells measured by non-imaging flow cytometry. D) Frequencies of DE cells detected by imaging flow cytometry using the optimal (OPT) imaging and the cell sorting matching (CSM) gating strategies, in comparison to cell sorting. Random galleries of DE cells identified by imaging flow cytometry with E) the OPT imaging gating and F) the CSM gating strategy. G) B cell and T cell gene scores derived from the scRNAseq data of sorted B cells, T cells and DE cells as reported in (5), and available under GEO accession number GSE129112. Data were derived from (A-C) 5 healthy subjects and (D-F) 3 healthy subjects.

When examining the scRNAseq data generated by Ahmed et al. (GEO accession number GSE129112), we found that DE cells have a significantly lower RNA concentration for any marker unique to either T cells or B cells (**Figure 4G**). Thus, the reported scRNAseq signature of sorted DE cells in Ahmed et al. is also consistent with cell-cell complexes.

## Discussion

In this study, we demonstrated that cell-cell complexes pairing a T cell with another immune cell type such as a monocyte or a B cell appear at first glance to be a distinct ‘halfway’ cell population with mixed lineage features at both protein and mRNA level. Further analyses identified key parameters from flow cytometry and scRNAseq data that are specific to cell complexes and thus indicate their presence within any cell population identified by flow cytometry. We also suggested experiments that can be used by researchers facing a case of suspicious dual-expressing cells, including imaging flow cytometry with appropriate gating strategy and direct microscopy analysis of sorted cells. Applying this knowledge to the DE cells identified in the study of Ahmed et al, we found that DE cells from healthy PBMC have a flow cytometry signature of doublets, which was also confirmed by imaging flow cytometry. Importantly, scRNAseq data from Ahmed et al also showed a DE signature of doublets. This observation emphasizes the importance of systematically and rigorously assessing the presence of cell-cell complexes in newly described cell populations, especially in cases where they display mixed lineage properties.

So why do we think that the Ahmed et al. paper might have overlooked the presence of complexes in their DE cell population expressing both T cell and B cell markers? The paper does show convincing evidence that single DE cells are present in imaging flow cytometry data. While the imaging data presented in the paper is limited to a single cell, the staining for both TCR and IgD is uniform on the cell surface, and it represents a convincing dual expresser. Thus, our concern is not that the DE cell presented in the Ahmed study is not real, but rather that large parts of their subsequent experiments were solely based on non-imaging flow cytometry and the assumption that DE cells are always singlets. Given the limitations of non-imaging flow cytometry to distinguish cell-cell complexes from singlet cells aforementioned, this is potentially problematic. Their gating strategy for imaging flow cytometry was described as ‘focused cells were selected on the basis of gradient RMS and an aspect ratio that was consistent with single cell events and devoid of debris or multi-cellular events (doublets)’ (5). Thus, it appears that the optimal imaging gating strategy was used here, which we have shown does not reflect adequately what is being detected by cell sorters. In this study, we have examined PBMCs from healthy donors and found that in non-perturbed settings, DE cells do have a flow cytometry signature of T cell-B cell complexes, with few dual expressing cells. The presence of dual expressers in the DE cell population could thus be unique to the three type I diabetes patients included in the Ahmed et al. study. However, the fact that sorted DE cells in the Ahmed study have a scRNA-seq signature of complexes suggest that our conclusion might also hold for the Ahmed et al. study as well. Besides, dual expressers identified by imaging flow cytometry might not necessarily represent a novel immune cell type, but could also be conventional T cells or B cells that have acquired the other lineage markers at the protein level following cell-cell interaction by trogocytosis, a common phenomenon occurring between immune cells (7, 8).

Overall, given the challenges of distinguishing true dual expresser single cells from cell-cell complexes in non-imaging flow cytometry and cell sorting, we think it is extremely important to include appropriate controls in future publications reporting on single cell immune profiling. The presence of cell-cell complexes signatures in flow cytometry and scRNAseq data should be systematically checked and reported. Additional experiments other than imaging flow cytometry should be performed such as direct microscopy analysis on sorted cells. Imaging cell sorters are being developed but have not reached yet the speed and sensitivity required for the detection of rare events, including cell-cell complexes. Until such technology is made commercially available, these recommendations will be crucial to ensure reliable interpretation of single cell datasets derived from flow cytometry, especially those reporting on the discovery of novel immune cell populations.

## Material and methods

### Subjects and samples

PBMC were obtained from healthy subjects by density gradient centrifugation (Ficoll-Hypaque, Amersham Biosciences) from leukapheresis or whole blood samples, according to the manufacturer’s instructions. Cells were resuspended to 10 to 50 million cells per mL in FBS (Gemini Bio-Products) containing 10% dimethyl sulfoxide (Sigma) and cryopreserved in liquid nitrogen. Cryopreserved PBMC were quickly thawed by incubating each cryovial at 37°C for 2 minutes, and cells transferred into 9 ml of cold medium (RPMI 1640 with L-Glutamin and 25 mM Hepes (Omega Scientific), supplemented with 5% human AB serum (GemCell), 1% Penicillin Streptomycin (Gibco) and 1% Glutamax (Gibco)) and 20 U/mL Benzonase Nuclease (Millipore) in a 15-ml conical tube. Cells were centrifuged and resuspended in medium to determine cell concentration and viability using Trypan blue and a hematocytometer and kept on ice until further analysis.

### Non-imaging flow cytometry

PBMC staining and acquisition were performed as described in (2). Briefly, cryopreserved PBMC were quickly thawed as described above, then cells (1-10 million) were transferred into a 15-ml conical tube, centrifuged, resuspended in 100 μl of PBS containing 10% FBS and incubated for 10 minutes at 4°C. Cells were then stained with 100 μl of PBS containing fixable viability dye eFluor506 (eBiosciences), 2 μl of Trustain FcR blocking reagent (BioLegend), 3 μl of anti-human CD45 antibody (clone HI100, eBiosciences) and antibody cocktail 1 or 2 for 20min at room temperature. Cocktail 1 was used to identify CD3+CD14+ cells and contained 3 μl of anti-human CD3-AF700 antibody (clone UCHT1, Biolegend) and 1 μl of anti-human CD14-PE antibody (clone 61D3, eBiosciences). Cocktail 2 was used to identify DE cells and contained 3 μl of anti-human CD5-APCeF780 antibody (clone L17F12, Biolegend), 2 μl of anti-human CD19-PECy7 antibody (clone HIB19, TONBO biosciences), and 3 μl of anti-human TCRab-AF488 antibody (clone IP26, Biolegend). After 2 washes in staining buffer (PBS containing 0.5% FBS and 2mM EDTA (pH 8.0), cells were resuspended into 100-500 μl of staining buffer, transferred into a 5 ml polypropylene FACS tube (BD Biosciences) and stored at 4°C protected from light for up to 4 hours until flow cytometry acquisition. Acquisition was performed on a BD FACSAria III cell sorter (BD Biosciences). Performance of the cell sorter was checked daily by the flow cytometry core at La Jolla Institute with the use of CS&T beads (BD Biosciences), and PMT voltages were manually adjusted for optimum fluorescence detection on each experiment day. Compensation was realized with single-stained beads (UltraComp eBeads, eBiosciences) in PBS using the same antibody volume as for the cell staining. Gating strategies are available on Figure 1A for T cells, monocytes and CD3+CD14+ cells and on Figure 4A for T cells, B cells and DE cells.

### Imaging flow cytometry

For the visualization of CD3+CD14+ cells, frozen PBMC were thawed and stained with 2 μl of anti-human CD3-AF488 (clone UCHT1, Biolegend) and 1 μl of anti-human CD14-PE (clone 61D3, eBiosciences) as described in the flow cytometry section above. For the visualization of DE cells, frozen PBMC were thawed and stained with 3 μl of anti-human CD5-APCeF780 antibody (clone L17F12, Biolegend), 2 μl of anti-human CD19-PECy7 antibody (clone HIB19, TONBO biosciences), and 3 μl of anti-human TCRab-AF488 antibody (clone IP26, Biolegend), as described in the flow cytometry section above. After two washes in PBS, cells were resuspended to 10×10^6^ cells/mL in staining buffer containing 5μg/mL Hoechst (Invitrogen) and 1μg/mL 7-AAD (Biolegend) and stored at 4°C protected from light until acquisition. Acquisition was performed with ImageStreamX MkII (Amnis) and INSPIRE software version 200.1.620.0 at 40X magnification and the lowest speed setting. A minimum of 4,000 CD3+CD14+ or CD5+CD19+ events in focus were collected. Compensation was performed using single stained cells. Data analysis was performed using IDEAS version 6.2.183.0.

### Cell sorting

PBMC were stained with fixable viability dye eFluor506 (eBiosciences), 3 μl of anti-human CD3-AF700 antibody (clone UCHT1, Biolegend) and 1 μl of anti-human CD14-PE antibody (clone 61D3, eBiosciences), as described in the flow cytometry section above. T cells, monocytes and CD3+CD14+cells were identified based on the gating strategy presented in Figure 1A. Sorts were performed on a BD Aria III cell sorter directly into a 96-well PCR plate containing 4 μl cell lysis buffer for single cell sorts, or in Eppendorf tubes containing staining buffer for bulk sorts.

### Single cell RNA sequencing

Single cell RNA sequencing was performed using Smart-seq2 as previously described (9). RNA loss was minimized by performing on-plate RNA capture, reverse-transcription and whole transcriptome pre-amplification (24 cycles) that results in ~1–30 ng of cDNA as previously described (9). 0.3 to 0.5 ng of pre-amplified cDNA was used to generate barcoded Illumina sequencing libraries (Nextera XT library preparation kit, Illumina) in an 8μL reaction volume. Multiple quality-control steps were included to ensure consistency during the procedure for all samples. Samples that failed at quality-control steps as described in (9, 10), were eliminated from downstream procedures and analysis. Libraries were pooled and sequenced on the HiSeq2500 Illumina platform to obtain more than 2 million 50-bp single end reads per cell. Single cell RNA sequencing data were mapped against the human hg19 reference genome using TopHat (v1.4.1., - -library-type fr-secondstrand -C) and Gencode version 19 (GRCh37.p13) as gene model reference for alignment. Sequencing read coverage per gene was counted using HTSeq-count (-m union -s yes -t exon -i gene_id, http://www.huber.embl.de/users/anders/HTSeq/). Counts per gene are obtained by counting all the transcripts mapping to a gene and are together referred to as transcript. To normalize for sequencing depth and varying transcript length counts were TPM (Transcripts Per Million reads) transformed. For PCA analysis only genes with at least 20 TPM were considered. PCA was performed in Python using SciPy SVD algorithm and plotted with Matplotlib (11). Cell-type specific scores were calculated by summing the TPM counts of 100 selected genes for each cell type. Monocytes genes were selected as the 100 first genes with the highest TPM counts in monocytes, and null expression in T cells. Similarly, T cells genes were selected as the 100 first genes with the highest TPM counts in monocytes, and null expression in monocytes. Raw and normalized data are available under accession number GSE117435. T cell and B cell scores were calculated in a similar fashion, using the data available under GEO accession number GSE129112.

### Microscopy

CD3-CD14+ monocytes, CD3+CD14-T cells and CD3+CD14+ cells were bulk sorted as described in the cell sorting section, and then transferred into individual chambers of a microscope slide. Fluorescence signals corresponding to CD3 and CD14 were detected with spectrally tuned PMTs using corresponding excitation lines of a Zeiss LSM 880 confocal microscope.

## Acknowledgments

We thank Dr. Cheryl Kim and all present and past members at the flow cytometry core facility at the La Jolla Institute for Immunology for assistance in cell sorting and technical discussion. We thank Dr. Zbigniew Mikulski from the microscopy core at the La Jolla Institute for Immunology for assistance and technical advice on microscopy imaging. We thank Yoav Altman at the Sanford Burnham Prebys flow cytometry core for technical assistance with imaging flow cytometry. Research reported in this manuscript was supported by the National Institute of Allergy and Infectious Diseases division of the National Institutes of Health under award numbers U19AI118626, S10OD021831 and S10OD016262, and by The Tullie and Rickey Families SPARK Awards at La Jolla Institute for Immunology. The content is solely the responsibility of the authors and does not necessarily represent the official views of the National Institutes of Health. Imaging flow cytometry was supported by the James B. Pendleton Charitable trust.

